# Evaluation of Bacteria Detected in Both Urine and Open Wounds in Nursing Facility Residents

**DOI:** 10.1101/614917

**Authors:** Josie Libertucci, Christine M. Bassis, Marco Cassone, Kristen Gibson, Bonnie Lansing, Lona Mody, Vincent B. Young, Jennifer Meddings

## Abstract

Nursing home residents are at a greater risk of developing pressure injuries that develop into an open wound. Open wounds can be colonized with bacteria from multiple sources. Emerging evidence suggests the specific composition of the open wound microbial community can result in delayed wound healing and increased infection risk. Understanding the factors that influence microbial colonization of open wounds, can lead to the prevention of infections. The relationship between bacteria found in urine and that of open wounds, is currently unknown. Recent studies have shown that nursing home residents can harbor a urinary microbiota independent of symptoms and frequently experience urinary incontinence. To determine if the colonization of open wounds mirrors urinary colonization, we conducted a pilot study with nursing home residents, comparing bacteria present in open pressure injuries below the umbilicus and urine. To identify microbial species that were present in both urine and open wound at one timepoint, standard bacteriologic culture techniques followed by MALDI-TOF was used as well as16S rRNA-encoding gene amplicon sequencing. In this study we found that some bacteria detected in urine were also detected in open wounds in one individual at one timepoint, using both culture dependent and independent techniques. Bacterial species that were more often detected at both sites included *Enterococcus faecalis, Proteus mirabilis, Escherichia coli*, and *Providencia stuartii*. This pilot study provides evidence that urinary colonization can mirror open wound colonization. Further studies are needed to investigate if parallel colonization of these anatomical sites affect infection outcomes.

## INTRODUCTION

Chronic open non-healing wounds severely impact a person’s quality of life and are estimated to impose $25 billion on the health-care system annually within the United States (1–3). Older adults are at greater risk of developing chronic open non-healing wounds, which can include pressure injuries (previously known as pressure ulcers), diabetic foot ulcers, arterial insufficiency ulcers, and venous leg ulcers (4–6). Pressure injuries in nursing home residents can occur related to skin breakdown over boney prominences related to immobility. It is important to focus on prevention of chronic wounds in this population, as the rate of wound repair and healing reduces with increased age (5). Additionally, bacterial colonization in open wounds can contribute to impaired wound healing (7, 8). Open pressure injuries in nursing home residents are not only susceptible to contamination of microorganisms from the environment, which includes contaminated caregivers, but also indigenous skin microorganisms can invade and become pathogenic once the skin barrier is damaged (9). These factors make it important to focus efforts on preventing bacterial contamination of open wounds. In order to design interventions to prevent wound contamination and invasive infection, it is imperative to understand all sources of possible contamination.

In older adults, urinary incontinence is common (8, 10, 11). When all other types of management and treatments for urinary incontinence fail to keep the skin dry, bladder catheterization is often performed. This is particularly important when the patient has an open wound in a body location, such as the sacrum, that is challenging to keep clear of urine or feces (10, 12–15). However, it is well known that indwelling urinary catheters can lead to increased bacterial colonization, including colonization of antibiotic resistant organisms, and can promote urinary tract infection (14, 16, 17). Within nursing home residents, urinary tract infections are the most common type of infection (18, 19). To prevent urinary tract infections, multiple recommendations regarding best approaches to catheter care have been made, including the reduced use of catheters in this patient population (20–23). Recent studies confirm the presence of a resident microbial community in urine from both catheterized and non-catheterized patients, however, the impact of these communities on wound contamination remains unknown (24–26).

In this pilot study, we sought to determine if bacteria found in urine of nursing home residents would also be found within open wounds. We hypothesized that open wounds located between the umbilicus and mid-thigh could harbor bacteria also found in the nursing home resident’s urine. To address this question, we identified bacteria from urine and open wounds using both culture dependent and independent methods to obtain a more complete census of the bacteria present at these two anatomical sites.

## MATERIALS AND METHODS

### Study design and participants

This study was approved by the institutional review board at the University of Michigan (HUM00092777). All participants (or approved decision makers) provided written informed consent prior to the initiation of this investigation. Subjects enrolled in this study were nursing home residents recruited from two nursing homes located in southeast Michigan. Recruitment for nursing home A occurred between March 2015 – May 2016 whereas recruitment from nursing home B occurred between January 2016 – May 2016. Both facilities included short and long-term care beds. In order for residents to be eligible for this study they had to be a minimum of 18 years of age and have an open wound in the anatomical area of interest (between umbilicus and mid-thigh). For the purposes of this investigation, open wounds included pressure injuries (at stages 2-4), and other open wounds with potential multiple etiologies (e.g., burns or surgical incisions) that were not anticipated to have a direct connection with the gastrointestinal or genitourinary tract. Nursing home residents with a closed wound (such as a stage 1 pressure injury) or wounds expected to have a connection with the gastrointestinal tract and/or urinary tract such as perirectal, fistula, and ostomies, were ineligible for this study. Additional criteria that deemed a resident ineligible to enroll in this study included: residents who were receiving end-of-life care or were anticipated to be discharged prior to the completion of the study, the presence of nephrostomy tube or ileal conduit urinary diversion as a primary means of urine output, or if the resident was anuric. Participants were subject to withdrawal from this study if their open wound healed, was discharged or admitted to the hospital, or retracted consent prior to the completion of sample collection. Patient data was collected by LM, JM, and BL and included: demographics, reason for admission, wound and catheter (if applicable) information, continence (fecal and urinary), functional status, mobility, relevant medical history, and comorbidities.

### Specimen Collection

All wound swabs were collected during regularly scheduled wound dressing changes by nursing staff in order to minimalize disturbance in the healing process. Some subjects were sampled at multiple timepoints. At each sampling timepoint, two wound swabs were taken using the Levine method. Briefly, wound swabs were taken by twirling the end of a sterile cotton-tipped applicator stick on the open wound in a one-square centimeter area for five seconds. Two types of cotton swabs were used to collect samples from open wounds –BactiSwab™ (Starplex Scientific, Inc, Ontario, Canada) and BactiSwab Dry™ (Remel, Lenexa, KS). BactiSwab™ was used to collect samples for downstream culture-dependent identification and BactiSwab Dry™ was used to collect samples for downstream culture-independent identification. Urine specimens were collected from residents at the same sampling timepoint as wound swabs were collected. Urine specimens were collected by midstream void (in residents able to void without a urinary catheter) and by aseptic collection from urinary catheter and aliquoted for culture-dependent and independent identification.

### Culture isolation and identification of bacterial species by MALDI-TOF analysis

Bacteria were isolated from wound swabs and urine samples by streaking specimens onto Trypticase Soy Agar with 5% Sheep Blood (PA-254053.07, Becton-Dickinson, Franklin lakes, NJ), Colombia Agar with 5% Sheep Blood (PA-254005.06, Becton-Dickinson, Franklin lakes, NJ), MacConkey Agar (L007388, Becton-Dickinson, Franklin lakes, NJ), and Bile Esculin Agar (dehydrated 299068, Becton-Dickinson, Franklin lakes, NJ). Plates were incubated at 35°C for 24 hours. Representative bacterial colonies were picked and grown in 5 ml of Brain Heart Infusion (BHI) media for 18 hours, shaking at 37°C (dehydrated DF0418-17-7, Becton-Dickinson, Franklin lakes, NJ). Freezer stocks were made by mixing 500 μl of liquid culture with 500 μl of glycerol (BP229-1, Fisher BioReagents, Pittsburgh, Pennsylvania) and stored at −80°C. Bacterial species were identified by streaking each freezer stock onto BHI agar (241830, Becton-Dickinson, Franklin lakes, NJ) using a 1 μl disposable inoculating loop (22-031-21, Fisher Scientific, Hampton, New Hampshire) and grown at 37°C for 24 hours. Colonies were identified using Matrix-Assisted Laser Desorption Ionization – Time of Flight (MALDI-TOF) analysis on a Bruker MALDI Biotyper CA System. One single colony was smeared, using the blunt end of a wooden toothpick, onto a reusable polished steel target plate (MSP 96 target polished steel BC 8280800, Bruker Daltonik, Bremen, Germany). For each MALDI-TOF run, two positive controls were added to the target plate, which included a Gram-negative control (*Escherichia coli*, ATCC 25922) and a Gram-positive control (*Staphylococcus aureus* ATCC 25923). The target plate was air-dried prior to the addition of a matrix solution and formic acid using the MALDI Biotyper Galaxy automated target plate preparation system (1836007 Bruker Daltonik, Bremen, Germany). Analysis was performed as per manufacturer’s instructions. Bacterial identification was taken from the identified best match with a score value ≥ 2.00.

### Identification of bacterial cultivars using Sanger sequence analysis of the 16S rRNA-encoding gene

Bacterial isolates were streaked onto BHI agar, from each freezer stock, using a 1 μl disposable inoculating loop and grown at 37°C for 24 hours. A single colony was grown in 5 ml of BHI media at 37°C for 18 hours, shaking. Bacteria were then harvested by centrifuging 1 ml of culture for 1 minute at 10,000 rpm. Harvested pellets were added to a MASTERBLOCK 96 deep well microplate (780271, VWR, Radnor, Pennsylvania) for genomic DNA extraction using the PowerMag Soil DNA isolation Kit (Mo Bio Laboratories, Inc., Cardlsbad, CA). Automated isolation was performed using an EpMotion 5075 instrument (Eppendorf, Hamburg, Germany). In some cases, genomic DNA was isolated by hand using the DNeasy Blood and Tissue Kit (69504, Qiagen, Venlo, Limburg, Netherlands). Molecular identification was completed by sequence analysis the 16S rRNA-encoding gene. One μl of extracted genomic DNA (1 − 20 ng/μl) was amplified in a 20 μl reaction that contained: 11.85 μl of ultra-pure DNase/RNase free distilled water (10977023, Invitrogen, Walthan, Massachusetts); 2 μl of 10x AccuPrime PCR Buffer II (12346-086, Invitrogen, Walthan, Massachusetts); 4 μM of forward primer (27F, 5’ AGA GTT TGA TCM TGG CTC AG 3’); 4 μM of reverse primer (1492R, 5’ GGT TAC CTT GTT ACG ACT T 3’); and 0.15 μl of *Taq* DNA polymerase (2346-086, Invitrogen, Walthan, Massachusetts). PCR amplification was performed under the following conditions: an initial denaturation step for 2 minutes at 95°C followed by 30 cycles of denaturation (95°C for 20 seconds), primer annealing (55°C for 15 seconds), and extension (72°C for 5 minutes), followed by a final extension step (72°C for 10 minutes). Amplicons were prepared for sequencing by adding 0.25 μl of ExoSAP-IT (78250, Affymetrix, Santa Clara, California) to 3.25 μl of ultra-pure DNase/RNase free distilled water and 1.5 μl of amplicon and incubated for 30 minutes at 37°C followed by 15 minutes at 80°C. Cleaned PCR products were sent for Sanger sequencing at the University of Michigan Medical School DNA Sequencing Core using nested 16S rRNA gene primers (338F, 5’ ACT CCT ACG GGA GGC AGC 3’ and 907R 5’ CCG TCA ATT CMT TTG AGT TT 3’). The forward and reverse reads were aligned and identification performed using the Ribosomal Database Project (RDP) classifier (Release 11, Update 5, September 30, 2016) (27).

### Urine and wound bacterial community analysis via 16S rRNA gene amplicon sequence analysis

For urine, 1 ml of urine was added to a glass bead tube (26000-50-BT, Mo Bio Laboratories Inc., Cardlsbad, CA) and centrifuged for 10 minutes at 5,000 x g. The supernatant was discarded, and this process was repeated using an additional 1 ml of urine. The urine pellets were then stored at −80°C until processed for DNA extraction. For wound samples, the tip of the wound swab was cut and placed into microcentrifuge tubes and stored at 80°C until processed. Genomic DNA from urine and wound swabs were extracted using the MAGATTRACT PowerMicrobiome DNA/RNA EP Kit (formerly known as the PowerMag Microbiome RNA/DNA Isolation Kit, 27500-4-EP, Mo Bio Laboratories Inc., Cardlsbad, CA) using the Eppendorf EpMotion liquid handling system.

The 16S rRNA-encoding gene from genomic DNA was amplified using barcoded dual-index primers developed by Kozich et al., 2013 (28). The process used for library generation has been previously described in Seekatz et al., 2015 (29). Barcoded dual-index primers specific to the V4 region of the 16S rRNA gene were used for PCR amplification. PCR reactions are composed of 5 μL of 4 μM equimolar primer set, 0.15 μL of AccuPrime Taq DNA High Fidelity Polymerase, 2 μL of 10x AccuPrime PCR Buffer II (Thermo Fisher Scientific, catalog no. 12346094), 11.85 μL of PCR-grade water, and 1 μL of DNA template. The PCR conditions used consisted of an initial denaturation 2 min at 95°C, followed by 30 cycles of 95°C for 20 s, 55°C for 15 s, and 72°C for 5 min, with a final extention at 72°C for 10 min. The DNA amplicons in each PCR reaction was normalized using the SequalPrep Normalization Plate Kit (Thermo Fisher Scientific, catalog no. A1051001). The normalized reactions were pooled and quantified using the Kapa Biosystems Library qPCR MasterMix (ROX Low) Quantification kit for Illumina platforms (catalog no. KK4873). The length of the pooled amplicons (~399 bp) was confirmedusing a high-sensitive DNA analysis kit (catalog no. 5067-4626) on an Agilent 3000 Bioanalyzer. Pooled amplicon library were sequenced on the Illumina MiSeq platform using the 500 cycle MiSeq V2 Reagent kit (catalog no. MS-102-2003) according to the manufacturer’s instructions with modifications of the primer set with custom read 1/read 2 and index primers added to the reagent cartridge. The “Preparing Libraries for Sequencing on the MiSeq” (part 15039740, Rev. D) protocol was used to prepare libraries with a final load concentration of 5.5 pM, spiked with 15% PhiX to create diversity within the run. DNA isolation and community sequencing was done by the University of Michigan Microbial Systems Molecular Biology Laboratory.

Sequence data was processed and analyzed using the software package mothur (v.1.39.5) and the MiSeq SOP (accessed on November 20, 2018) (28). Sequences were aligned to a recreated V4 specific SILVA SEED reference (release 119) and trimmed (30). Chimeras were then removed using uchime (31). Prior to calculating observed operational taxonomic units (OTUs), this dataset was normalized to 7,395 sequences per sample. OTUs were then created by clustering sequences based on 97% sequence similarity by the average neighbor method. OTU concordance between each urine and open wound sample per timepoint was calculated using the mothur calculator sharedsobs.

To identify the OTU classification for each bacterial cultivar, the following steps were taken. First, the full length 16S sequence was generated using Sanger sequencing as described above. Then, each sequence was aligned to the recreated V4 specific SILVA SEED reference created for the microbiome analysis as described. Once, the V4 sequence for each cultivar was identified, those sequences were aligned to the created OTUs. The output was matched to specific OTUs from the community analysis.

### Availability of data

Raw FASTQ files, including those for negative controls, were deposited in the SRA database under BioProject ID nunmber PRJNA533783. Detailed processing steps can be found in supplemental methods.

## RESULTS

### Characteristics of participants involved in this pilot study

In this pilot study, a total of 13 residents from two different nursing home facilities were recruited however, samples from only 9 residents were included in this study (Table 1). Four residents and their samples were excluded from this analysis for not having concurrent urine and wound samples. A total of 15 concurrent urine and wound samplings from 9 patients were included in the analysis with 6 subjects sampled at only 1 time point and 3 subjects sampled at multiple timepoints. The mean age for residents included in this analysis was 70 years old and all residents were male and Caucasian. Residents enrolled in this study remained in their respective nursing home for a mean of 131 days, with a range of 10-740 days. Most residents in this study were described as immobile (n=8, 89%). Of those 8 residents, immobility was described to be resulted from paraplegia (n = 4), morbid obesity (n = 2), and unknown (n=2). A proportion of residents in this study were documented as having urinary incontinence (n=4, 44%), two of which also had fecal incontinence. Each resident, with the exception of resident number 8, had only one eligible open wound in the area of interest. Resident 8 had two eligible open wounds in the area of interest.

**Table 1:** Participant characteristics at first sampling timepoint

| <u>Characteristic</u> | <u>Residents (n=9)</u> |
| --- | --- |
| **Demographics** |  |
| Age, mean | 70 |
| Male Sex, no. (%) | 9 (100%) |
| Race, no. (%)<br>Caucasian | 9 (100%) |
| Length of stay days, mean [range] | 131 [10-740] |
| **Mobility** |  |
| Immobile | 8 (89%) |
| Paraplegia | 4 (44%) |
| Morbid Obesity | 2 (22%) |
| Etiology not further specified | 2 (22%) |
| **Incontinence** |  |
| Incontinent | 4 (44%) |
| Fecal and Urinary Incontinence | 2 (22%) |
| Urinary (without Fecal) Incontinence | 2 (22%) |
| **Method for Urine Specimen Collection** |  |
| Clean Catch (no catheter use) | 3 (33%) |
| Intermittent Straight Catheter (ISC) | 4 (44%) |
| Indwelling Transurethral (Foley) Catheter | 2 (22%) |

### Microbial community structure of urine and open wounds

To obtain a global view of bacteria present in urine and wound from these residents, we used culture-independent analysis that involved sequence analysis of V4 region amplicons of the 16S rRNA gene. This method has been used to characterize the community structure of complex microbial consortia from a variety of habitats. A total of 1,411,374 sequences were generated in this dataset with an average of 45,528 sequences per sample (min = 7,395 max = 141,600 sequences per sample). This analysis revealed the presence of a multispecies bacterial community in most samples (Figure 1). In general, a greater diversity of organisms was present in wound samples compared to urine samples. Urine samples had a mean observed species (richness) of 44.6 (min = 5; max = 151; standard deviation = 39.96), compared to an average of 74.5 for wounds samples (min = 19; max = 164; standard deviation = 49) (Supplemental Table 1). Residents varied with respect to the nature of the community structure in both urine and open wounds. In residents who had more than one sample obtained over time, the communities at one site were more similar over time compared to the communities in other residents.

**Figure 1:**
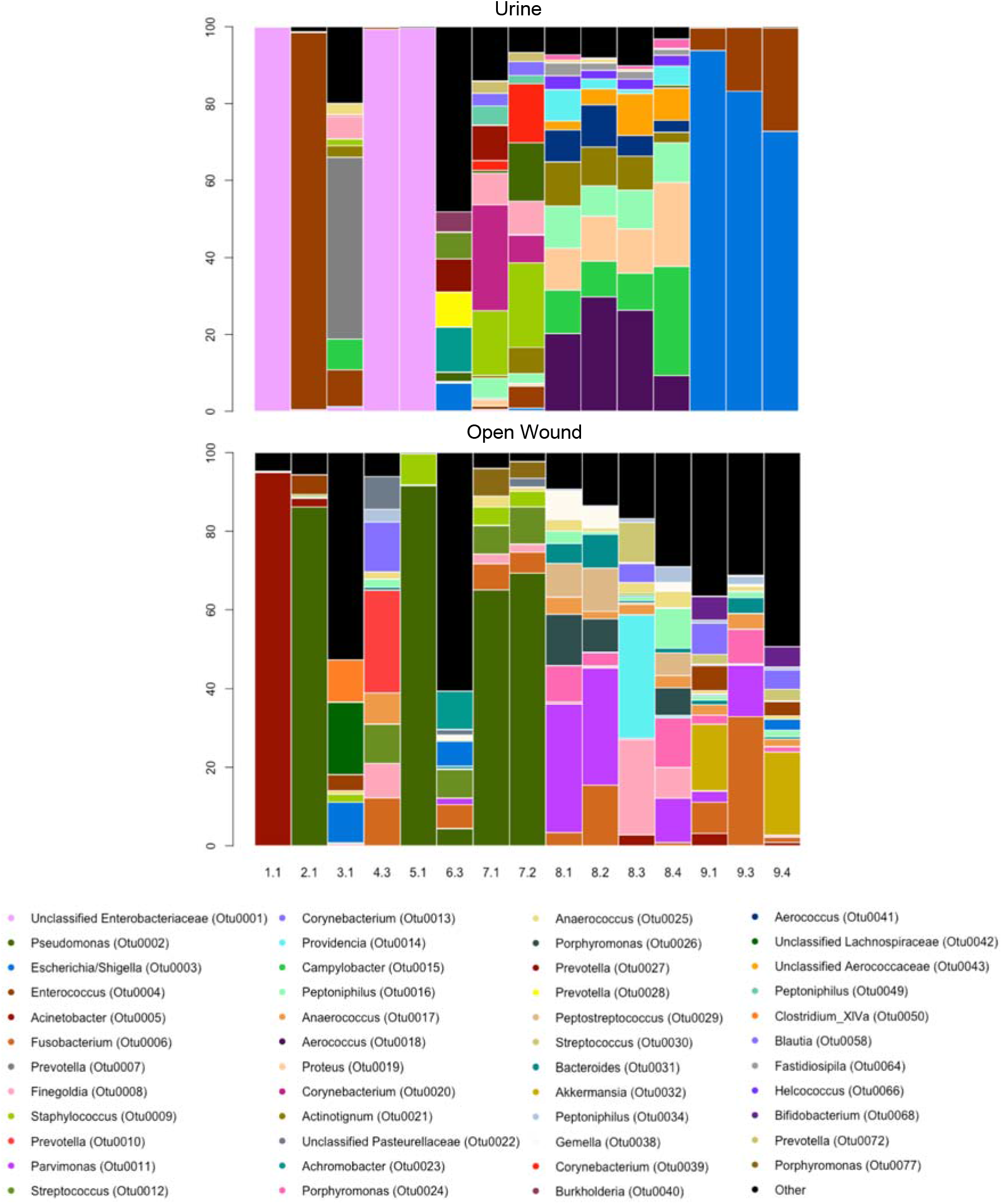
Microbial community structure of urine and open wounds in this study. The microbial community structure of urine and open wounds was determined through amplification of the V4 16S rRNA gene followed by Illumina sequencing. Sequence data was processed and analyzed using the software package mothur. Samples were normalized to 7,395 sequences per sample and barplot was visualized using R. In some cases, urine samples were dominated by one OTU, whereas wound samples were more frequently found to be characterized by many species. The X-axis identifies the resident and sampling time point (e.g., 1.1 for resident 1 at the first sampling timepoint). The y-axis represents the relative abundance for each OTU.

In addition to the greater diversity observed in wound samples compared to urine, urine was more likely to be characterized by the presence of one dominant OTU compared to wounds (Figure 1). For example, OTU0001 (Unclassified Enterobacteriaceae) was the only type of bacteria seen in urine samples from residents 1, 4 and 5 while the three urine samples from resident 9 were dominated by OTU0003 (*Escherichia/Shigella*). In wound samples, greater diversity was observed, although a minority of communities had one dominant OTU. For example, resident 1 had an overabundance of OTU0005 (*Acinetobacter*), whereas residents 2, 5, and 7 had an overabundance of OTU0002 (*Pseudomonas*).

### Culture-based analysis of urine and wound specimens

For clinical purposes, bacterial culture has been the mainstay of microbial isolation and identification. We isolated bacteria from urine and open wounds using aerobic culture on standard microbiologic media and identified organisms obtained through the use of MALDI-TOF (Supplemental Table 2). One hundred and eighteen bacterial isolates were obtained from 15 wound and 15 urine samples. These 118 isolates were identified as a total of 31 different species. In 12 of 15 urine samples and 15 of 15 wound samples, more than one species was identified by culture. Within urine samples, the most frequently isolated bacteria were *Enterococcus faecalis, Aerococcus sanguinicola, Escherichia coli, Proteus mirabilis*, and *Providencia stuartii*. Within open wounds, *Staphylococcus* and *Corynebacterium* species were commonly isolated from residents, as well as *E. faecalis, Pseudomonas aeruginosa*, and *Acinetobacter baumannii*.

To compare the results of cultivation to the culture-independent analysis, partial 16S rRNA-encoding gene sequences were obtained from each cultivar. These partial 16S sequences were compared to the community 16S data to assign each cultivar to an OTU identified by the culture-independent analysis. With two exceptions (both were cultivars identified as *Corynebacterium aurimucosum* by MALDI-TOF), all of the cultivars mapped to an OTU identified by community analysis (Supplemental Table 2). For 6 of 15 urine samples, the cultivated organisms were the most abundant OTUs encountered by community analysis. This was comparable to open wound samples where for 5 out of 15 samples, the cultivar was the most abundant OTU (Table 2).

**Table 2:**
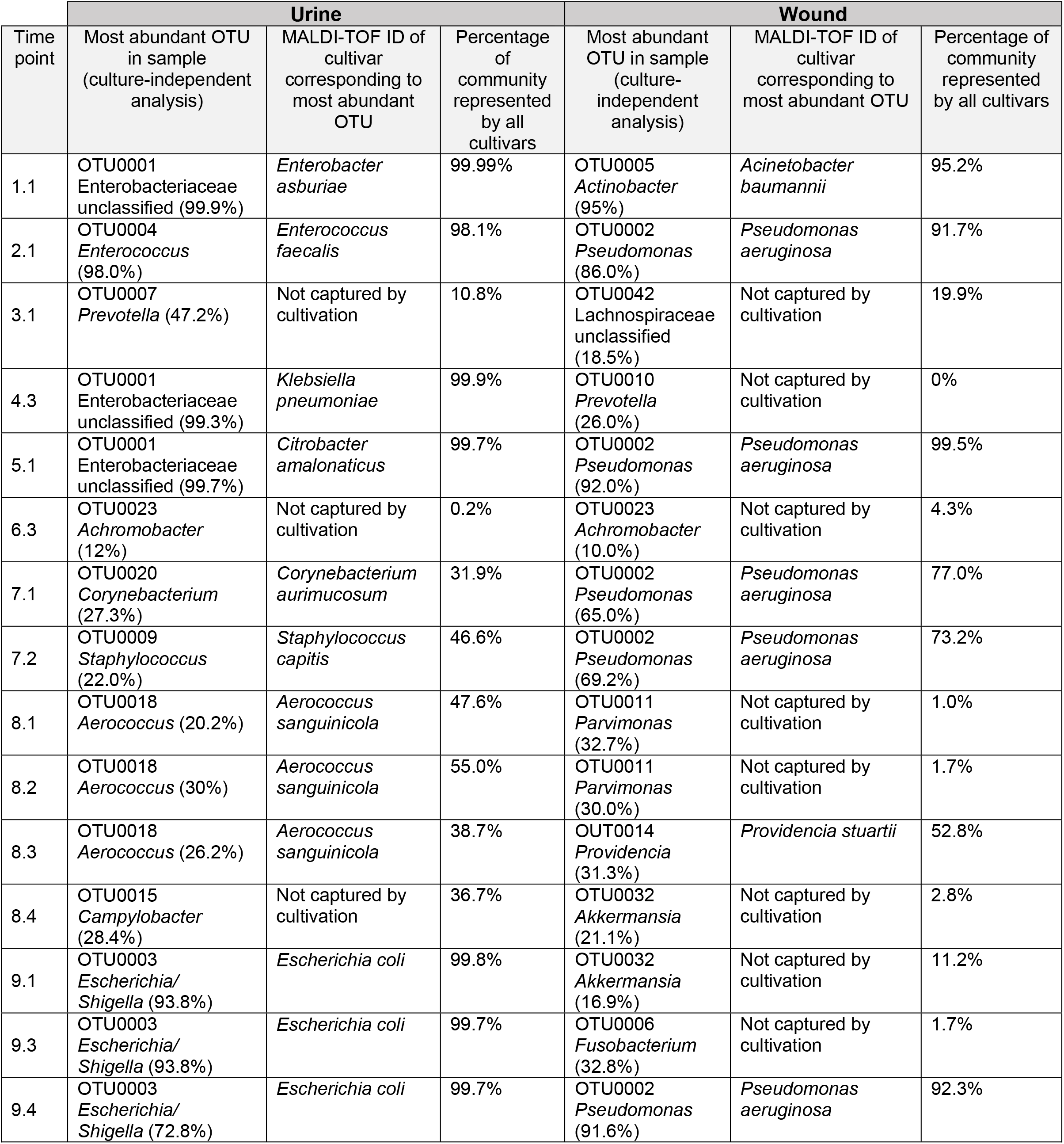
Abundance of cultivar identified OTUs within each sample

| Time point | Urine |  |  | Wound |  |  |
| --- | --- | --- | --- | --- | --- | --- |
|  | Most abundant OTU in sample (culture-independent analysis) | MALDI-TOF ID of cultivar corresponding to most abundant OTU | Percentage of community represented by all cultivars | Most abundant OTU in sample (culture-independent analysis) | MALDI-TOF ID of cultivar corresponding to most abundant OTU | Percentage of community represented by all cultivars |
| 1.1 | OTU0001<br>Enterobacteriaceae unclassified (99.9%) | *Enterobacter asburiae* | 99.99% | OTU0005<br>*Actinobacter* (95%) | *Acinetobacter baumannii* | 95.2% |
| 2.1 | OTU0004<br>*Enterococcus* (98.0%) | *Enterococcus faecalis* | 98.1% | OTU0002<br>*Pseudomonas* (86.0%) | *Pseudomonas aeruginosa* | 91.7% |
| 3.1 | OTU0007<br>*Prevotella* (47.2%) | Not captured by cultivation | 10.8% | OTU0042<br>Lachnospiraceae unclassified (18.5%) | Not captured by cultivation | 19.9% |
| 4.3 | OTU0001<br>Enterobacteriaceae unclassified (99.3%) | *Klebsiella pneumoniae* | 99.9% | OTU0010<br>*Prevotella* (26.0%) | Not captured by cultivation | 0% |
| 5.1 | OTU0001<br>Enterobacteriaceae unclassified (99.7%) | *Citrobacter amalonaticus* | 99.7% | OTU0002<br>*Pseudomonas* (92.0%) | *Pseudomonas aeruginosa* | 99.5% |
| 6.3 | OTU0023<br>*Achromobacter* (12%) | Not captured by cultivation | 0.2% | OTU0023<br>*Achromobacter* (10.0%) | Not captured by cultivation | 4.3% |
| 7.1 | OTU0020<br>*Corynebacterium* (27.3%) | *Corynebacterium aurimucosum* | 31.9% | OTU0002<br>*Pseudomonas* (65.0%) | *Pseudomonas aeruginosa* | 77.0% |
| 7.2 | OTU0009<br>*Staphylococcus* (22.0%) | *Staphylococcus capitis* | 46.6% | OTU0002<br>*Pseudomonas* (69.2%) | *Pseudomonas aeruginosa* | 73.2% |
| 8.1 | OTU0018<br>*Aerococcus* (20.2%) | *Aerococcus sanguinicola* | 47.6% | OTU0011<br>*Parvimonas* (32.7%) | Not captured by cultivation | 1.0% |
| 8.2 | OTU0018<br>*Aerococcus* (30%) | *Aerococcus sanguinicola* | 55.0% | OTU0011<br>*Parvimonas* (30.0%) | Not captured by cultivation | 1.7% |
| 8.3 | OTU0018<br>*Aerococcus* (26.2%) | *Aerococcus sanguinicola* | 38.7% | OUT0014<br>*Providencia* (31.3%) | *Providencia stuartii* | 52.8% |
| 8.4 | OTU0015<br>*Campylobacter* (28.4%) | Not captured by cultivation | 36.7% | OTU0032<br>*Akkermansia* (21.1%) | Not captured by cultivation | 2.8% |
| 9.1 | OTU0003<br>*Escherichia/Shigella* (93.8%) | *Escherichia coli* | 99.8% | OTU0032<br>*Akkermansia* (16.9%) | Not captured by cultivation | 11.2% |
| 9.3 | OTU0003<br>*Escherichia/Shigella* (93.8%) | *Escherichia coli* | 99.7% | OTU0006<br>*Fusobacterium* (32.8%) | Not captured by cultivation | 1.7% |
| 9.4 | OTU0003<br>*Escherichia/Shigella* (72.8%) | *Escherichia coli* | 99.7% | OTU0002<br>*Pseudomonas* (91.6%) | *Pseudomonas aeruginosa* | 92.3% |

### Bacteria detected in both urine and open wounds per resident

We used both culture-dependent and independent methods to identify the presence of bacterial species within both urine and open wounds obtained at the same time (Table 3). Using bacterial culture, at least one bacterial species was detected in both urine and open wound for 13 out of 15 timepoints. In two cases, all bacterial species that were detected in urine were detected within a resident’s open wound (resident 3 sampling timepoint 1, and resident 7 sampling timepoint 2). Using culture-independent characterization of the wound and urine microbiota, at least 1 OUT was shared by both urine and open wounds for all 15 sampling timepoint.

**Table 3:** Number of bacterial species or OTUs detected both in urine and open wounds at the same sampling timepoint per resident

| Resident | Sampling timepoint | <u>Culture Dependant</u> |  |  | <u>Culture Independent</u> |  |  |
| --- | --- | --- | --- | --- | --- | --- | --- |
|  |  | Urine (No. species detected) | Open wound (No. species detected) | Number of species found in both urine and wound at the same sampling timepoint | Urine (No. OTUs detected) | Open wound (No. OTUs detected) | Number of OTUs found in both urine and wound at the same sampling timepoint |
| 1 | 1 | 1 | 2 | 0 | 5 | 32 | 1 |
| 2 | 1 | 1 | 3 | 1 | 26 | 43 | 8 |
| 3 | 1 | 3 | 3 | 3 | 73 | 79 | 50 |
| 4 | 3 | 2 | 4 | 1 | 9 | 38 | 1 |
| 5 | 1 | 3 | 4 | 2 | 5 | 19 | 2 |
| 6 | 3 | 1 | 2 | 0 | 151 | 163 | 106 |
| 7 | 1 | 5 | 4 | 1 | 57 | 31 | 13 |
|  | 2 | 4 | 4 | 2 | 51 | 29 | 9 |
| 8 | 1 | 4 | 9 | 2 | 63 | 56 | 19 |
|  | 2 | 4 | 10 | 2 | 65 | 61 | 19 |
|  | 3 | 3 | 6 | 2 | 71 | 75 | 20 |
|  | 4 | 4 | 14 | 3 | 62 | 71 | 19 |
| 9 | 1 | 2 | 5 | 2 | 11 | 142 | 5 |
|  | 3 | 2 | 5 | 2 | 10 | 115 | 4 |
|  | 4 | 2 | 7 | 2 | 10 | 164 | 4 |

Of the 30 bacterial species isolated in at least one sample of urine or wound using culture-based analysis, there were 6 bacterial species isolated from both urine and open wound at the same sampling timepoint for a resident (Table 4). This included (in descending order of frequency) *E. faecalis, P. mirabilis, E. coli, P. stuartii, P. aeruginosa* and *Citrobacter farmeri*. Using culture-independent methods (Supplemental Table 3), the community analysis data revealed that 164 OTUs were shared between urine and open wounds. The OTUs that were most frequently found in both the urine and wound at the same sampling timepoint were OTU0003 (*Escherichia/Shigella*), OTU0004 (*Enterococcus*), OTU0031 (*Bacteroides*), OTU0017 (*Anaerococcus*), OTU0021 (*Actinotignum*), and OTU0025 (*Anaerococcus*).

**Table 4:**
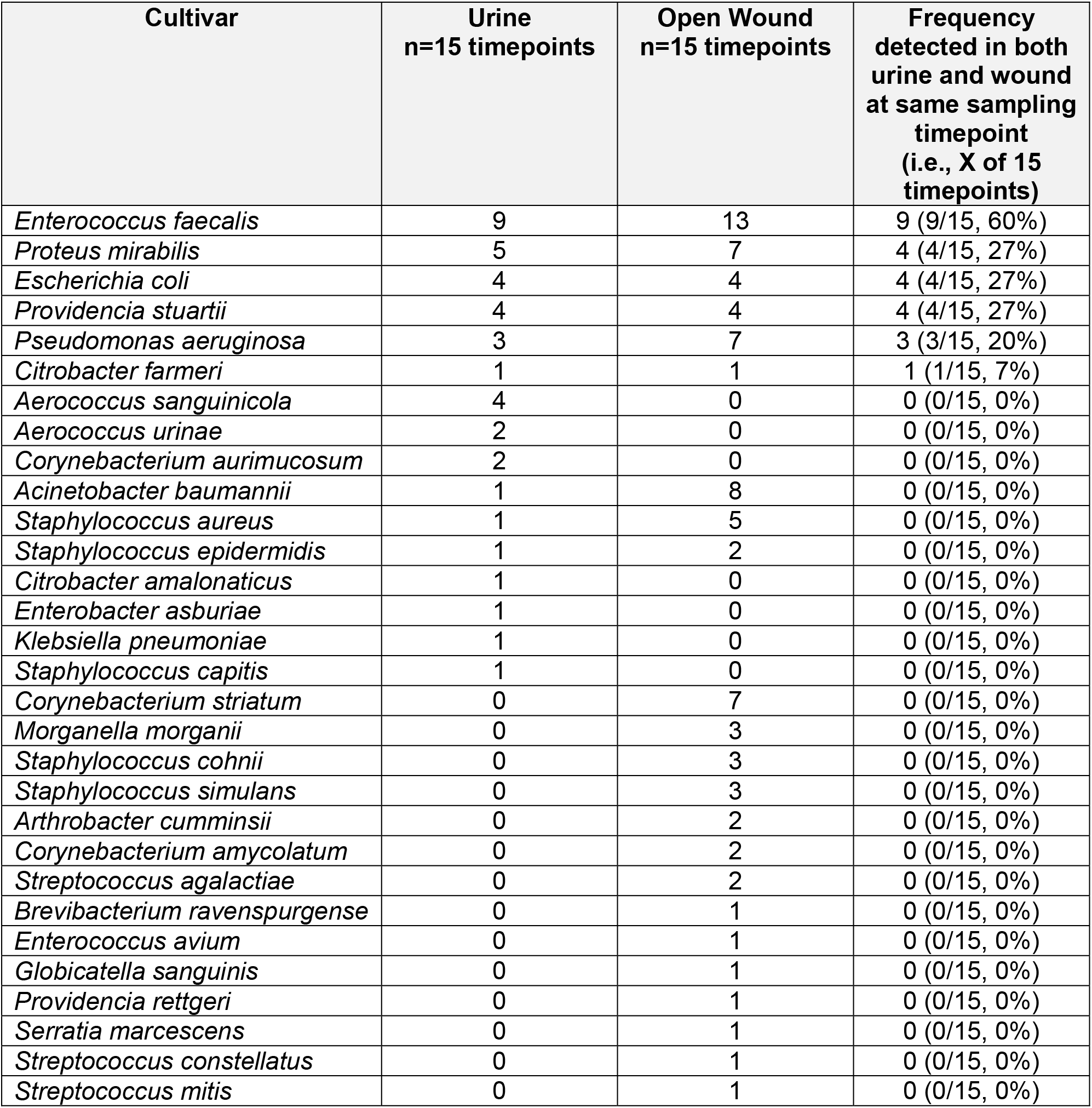
Frequency of bacterial species by culture-dependent identification in urine, open wound, and both urine and wound

## DISCUSSION

In this pilot study, we assessed how often the same bacterial species was identified in both the urine and open wounds from nursing home residents for specimens collected at the same sampling timepoint. This was assessed for wounds located in body areas anticipated to be at high risk of contamination from urine. Comparison of bacterial species identified from different anatomical sources in the same patient has been evaluated previously in an effort to identify possible sources of bacteremia (32), noting some organisms responsible for bloodstream infections were also found in urine and wound microbiota. Additionally, concordance between urinary tract infections and positive blood cultures in neonates has been previously identified (33). Although it is well recognized that contamination of open wounds by microorganisms can lead to infectious complications such as cellulitis, osteomyelitis, bacteremia, and delayed wound healing (5, 7, 34, 35), and that urine is frequently colonized (i.e., asymptomatic bacteriuria) in nursing home residents, we believe this is the first study to identify bacteria present in both urine and open wounds at the same timepoint, using both culture-dependent and culture-independent methods (36).

In this pilot study we identified bacterial species that were present in both urine and open wounds in a patient at a specific timepoint (Table 3). This finding was noted using both culture-dependent and culture-independent methods. Using culture dependent methods revealed that 13 out of 15 timepoints indicated at least one bacterial species was present in both urine and open wounds in one patient at a specific time. This is in contrast to culture-independent methods that revealed OTUs were shared at all timepoints sampled. Interestingly, there were some bacterial species that were found to be present in both urine and open wounds more often than other bacterial species. These species included *E. faecalis, P. mirabilis, E. coli, P. stuartii, P. aeruginosa*, and *C. farmeri*. This result was mirrored in our culture-independent analysis where some OTUs were also found to be shared more often than others. OTUs that were found to be present in both anatomical sites include OTU0003 (*Escherichia/Shigella*), OTU0004 (*Enterococcus*), OTU0031 (*Bacteroides*), OTU0017 (*Anaerococcus*), and OTU0021 (*Actinotignum*) (Supplemental Table 3). Overall these results indicate that bacterial colonization of the urine can mirror colonization of open wounds.

We initially performed this study by employing standard microbiologic culture to isolate organisms, and MALDI-TOF to identify those organisms, from both urine and open wounds. Since standard culture methods used for diagnosis and clinical treatment are designed to selectively enrich for target organisms, we sought to determine if the use of a more comprehensive, less biased method to retrieve information on the bacterial communities in urine and wounds would provide additional insight into the microbiota in these two anatomic sites. As expected, profiling the bacteria in urine and wounds using 16S rRNA-encoding gene sequence analysis provided evidence for a greater diversity of organisms than that encountered using culture alone (Supplemental Table 1). Organisms that were identified as the most abundant pathogen using culture were a minor component of the entire community in many samples based on culture-independent analysis. For example, in resident 3 at sampling timepoint 1, culture dependent identification revealed that *E. faecalis*, *E. coli*, and *C. farmeri* were the only species present in this urine specimen (Supplemental Table 2). When comparing these data to 16S rRNA-encoding gene sequence analysis, those species compose only 10.8% of the community, and the most abundant organism was found to be OTU0007 Prevotella (Table 2). In some cases, culture-independent analyses identified anaerobes as the most abundant OTUs identified in a given sample. This has been noted in previous studies that identified the presence of 16S genes from anaerobes in chronic wounds (37). It should be noted that identification of anaerobes from a sample using 16S rRNA-encoding gene sequence analysis does not indicate the presence of viable organisms. It is possible that this is simply a marker of fecal contamination.

This pilot study does have limitations. Shared bacterial species in both the wound and urine does not necessarily implicate the urine as the ultimate source of the microbes found in wounds. Indeed, with mostly cross-sectional sampling, the directionality of transfer itself is not known. Furthermore, we did not collect fecal samples, another potential source for contamination of both urine and wound. A limitation of our analysis is we did not collect fecal specimens from the residents to compare with urine and wound specimens. Indeed, some of the bacteria that were identified as most often being concordant between urine and wound including *E. faecalis*, *P. mirabilis*, *E. coli*, and *P. stuartii* can be found as indigenous members of the gut microbiota. This observation was also observed using culture-independent identification methods where we identified OTU0003 (*Escherichia/Shigella*), OTU0004 (*Enterococcus*), OTU0031 (*Bacteroides*), OTU0017 (*Anaerococcus*), OTU0021 (*Actinotignum*), and OUT0025 (*Anaerococcus*) were the species that were most concordant.

Overall, this pilot study indicates that bacteria from the urine can often be found in open wounds within the same patient at a point in time in nursing home residents, with or without urinary catheter use, using both culture dependent and independent methods. This study warrants further investigation into the role of urinary microorganisms within open wound contamination. Another important consideration is that some species are found at higher frequency than others, which suggest that they are more robust and can live in many environments. This is of particular concern as many of these organisms can often be found to be resistant to antibiotics and in some cases resistant to multiple types of antibiotics. Further investigation is needed to fully understand the relationship and clinical implications of organisms found in both urine and open wounds in the nursing home population.

